# Single-molecule sequencing of the *Drosophila serrata* genome

**DOI:** 10.1101/090969

**Authors:** Scott L. Allen, Emily K. Delaney, Artyom Kopp, Stephen F. Chenoweth

## Abstract

Long read sequencing technology promises to greatly enhance *de novo* assembly of genomes for non-model species. While error rates have been a large stumbling block, sequencing at high coverage allows reads to be self-corrected. Here we sequence and *de novo* assemble the genome of *Drosophila serrata*, a non-model species from the *montium* subgroup that has been well studied for clines and sexual selection. Using 11 PacBio SMRT cells, we generated 12 Gbp of raw sequence data comprising approximately 65x whole genome coverage. Read lengths averaged 8,940 bp (NRead50 12,200) with the longest read at 53 Kbp. We self-corrected reads using the PBDagCon algorithm and assembled the genome using the MHAP algorithm within the PBcR assembler. Total genome length was 198 Mbp with an N50 just under 1 Mbp. Contigs displayed a high degree of arm-level conservation with *D. melanogaster*. We also provide an initial annotation for this genome using *in silico* gene predictions that were supported by RNA-seq data.

## INTRODUCTION

Second-generation sequencing (2GS) platforms, such as Illumina sequencing-by-synthesis, have dramatically reduced genome sequencing costs while increasing throughput exponentially (Shendure and Ji 2008). The relatively low cost and massive throughput of second-generation sequencing platforms have paved the way for sequencing and *de novo* assembly of thousands of species’ genomes (Alkan et al. 2011). Second-generation sequencing methods generate short reads (less than a few hundred base pairs in length) that have limitations for *de novo* genome assembly, where assembly is performed without the aid of a reference genome (Green 1997; Miller et al. 2008; Nagarajan and Pop 2013; Alkan et al. 2011). With short reads, *de novo* assembly is an inherently difficult computational problem because repetitive DNA sequences are often much longer than the length of each read (Ukkonen 1992). For instance, it has been estimated that short read *de novo* assemblies could be missing up to 20% of sequence information because repeat DNA sequences can increase the number of misassembled and fragmented regions (Schatz et al. 2010; Alkan et al. 2011; Ukkonen 1992). One way to alleviate the problem of repetitive DNA in the *de novo* assembly process has been to incorporate a second set of mate-pair libraries with very long inserts (>2 kbp) (Li et al. 2010; Chaisson et al. 2009; Simpson et al. 2009; Alkan et al. 2011; Butler et al. 2008). Mate-pair libraries can resolve repeats (Treangen and Salzberg 2012; Wetzel et al. 2011) and improve scaffolding (van Heesch et al. 2013), but paired-end contamination and insert size mis-estimation can also lead to mis-assemblies (Phillippy et al. 2008; Sahlin et al. 2016).

More recently, third-generation (3GS) single-molecule sequencing technologies such as Pacific Biosciences’ (PacBio) SMRT sequencing and Oxford Nanopore’s MinlON sequencing, which currently produce much longer reads, up to 54 kbp (Lee et al. 2014) and > 10 kbp (Quick et al. 2014), respectively, can overcome some of the shortcomings of 2GS assembly (Berlin et al. 2015). Although long-read sequencing technology produces reads with a high error rate, ranging from 82.1% (Chin et al. 2011) to 84.6% accuracy (Rasko et al. 2011), sequencing errors occur at more or less random positions across long-reads (Chin et al. 2013) and can be corrected with 2GS short-read data (Koren et al. 2012) or by using excess 3GS reads for a self-correction (Chin et al. 2013).

In this paper, we use PacBio long-read sequencing to *de novo* assemble the genome of the fly, *Drosophila serrata,* which has been particularly well studied from an evolutionary standpoint. *D. serrata* is a member of the *Drosophila montium* subgroup, which split from the *D. melanogaster* subgroup approximately 40 Mya (Tamura et al. 2004), and consists of an estimated 98 species (Brake and Bächli 2008). At present, only one draft genome assembly *(D. kikkawai)* is available (Chen et al. 2014) from this species-rich subgroup. *D. serrata* has a broad geographical distribution, ranging from Papua New Guinea to south eastern Australia and has emerged as a powerful model for addressing evolutionary questions such as the evolution of species borders (Blows and Hoffman 1993; Hallas et al. 2002; Magiafoglou et al. 2002) and climate adaptation (Frentiu and Chenoweth 2010; Kellermann et al. 2009). The species has also been used to investigate sexual selection (Hine et al. 2002; Chenoweth et al. 2015), male mate choice (Chenoweth and Blows 2003; Chenoweth et al. 2007), mate recognition (Higgie et al. 2000), sexual dimorphism (Chenoweth et al. 2008; Yassin et al. 2016), sexual conflict (Delcourt et al. 2009) and indirect genetic effects (Chenoweth et al. 2010b). Its cuticular hydrocarbons, which serve as contact pheromones (Chung et al. 2014), have been extensively used to develop novel multivariate quantitative genetic approaches for exploring genetic constraints on adaptation (Blows et al. 2004; Chenoweth et al. 2010a; McGuigan et al. 2011b; Rundle et al. 2009).

Despite the importance of *D. serrata* as a model for evolutionary research, our poor understanding of its genome remains a significant limitation. Linkage and physical genome maps are available (Stocker et al. 2012), and an expressed sequence tag (EST) library has been developed (Frentiu et al. 2009), but the species lacks a draft genome. Here we report the sequencing and assembly of the *D. serrata* genome using exclusively Pacific Biosciences SMRT technology. We also provide an initial annotation of the genome based on *in silco* gene predictors and mRNA-seq data. Our *de novo* genome and its annotation will provide a resource for ongoing population genomic and trait mapping studies in this species as well as facilitate broader studies of genome evolution in the family Drosophilidae.

## MATERIALS AND METHODS

### Fly Strains and DNA Extraction

We sequenced a mix of ~100 mg of males and females from a single inbred line that originated from Forster, Australia, and had been inbred via full-sib mating for 10 generations before being maintained at a large population size (N ~ 250 individuals) (McGuigan et al. 2011b). A single further generation of full-sib inbreeding was applied before extraction of DNA. This same inbred line was used for the *D. serrata* linkage map, was the founding line for previous mutation accumulation studies (Latimer et al. 2015; McGuigan et al. 2014a; McGuigan et al. 2014b; McGuigan et al. 2011a) and is fixed for the light female abdominal pigmentation phenotype mapped by Yassin et al. (2016). High molecular weight DNA was extracted from fly bodies (heads were excluded to reduce eye pigment contamination) using a Qiagen Gentra Puregene Tissue Kit (Cat #158667) which produced fragments > 100 kbp (measured using pulsed-field gel electrophoresis). Two phenol-chloroform extractions were performed by the University of California-Davis DNA Technologies Core prior to preparation of a sequencing library.

### Genome Sequencing and Assembly

DNA was sequenced using 11 SMRT cells and P6-C4 chemistry on the Pacific Biosciences RS II platform. In total this produced ~13 billion base pairs spanning 136,119 filtered subreads with a mean read length of 8,840 bp and an N50 of 12,220 bp (Figure S1). The PacBio genome was assembled using the PBcR pipeline that implements the MHAP algorithm within the Celera Assembler (Berlin et al. 2015) and polished with Quiver (Chin et al. 2013) in three steps: (1) errors were corrected in reads using PBDagCon, which requires at least 50x genome coverage and utilizes the consensus of over-sampled sequences (Chin et al. 2013), (2) overlapping sequences were assembled using MHAP and the Celera Assembler (Berlin et al. 2015), and (3) contigs were polished with Quiver to correct for spurious SNP calls and small indels (Chin et al. 2013). The “sensitive” setting was used for both read correction and genome assembly (Berlin et al. 2015) whereas the default settings were used for polishing with Quiver (Chin et al. 2013). We elected to correct all reads as opposed to the default longest 40x. The longest 25x corrected reads were subsequently used for genome assembly. The PBDagCon correction was performed on a computer with 60 CPU cores and 1TB of RAM; 58 CPU cores were used for the assembly and the amount of RAM used, although not tracked, was far less than machine capacity. Error correction with PBDagCon took ~26 days. Assembly of corrected reads using MHAP and the Celera Assembler took ~19 hours using 28 CPU cores. Our initial runs using the much faster error correction algorithm (HGAP) produced a slightly shorter assembly (194 Mbp compared to 198 Mbp) with a slightly lower N50 (0.88 Mbp vs 0.95 Mbp). We therefore chose to use the more sensitive PBDagCon correction method.

### Transcriptome Sequencing and Assembly

The same inbred fly strain that was the progenitor for DNA sequencing was also used for adult mRNA sequencing to annotate the *D. serrata* genome. Adult males and females were transferred to fresh vials shortly after eclosion and held in groups of ~25 where they were allowed to mate and lay eggs for 2 days. They were then sexed under light CO_2_ anesthesia and snap frozen using liquid nitrogen in groups of 10, at the time of freezing all flies were assumed to be non-virgins. Total RNA was extracted from each pool of flies using the standard Trizol protocol. Initial quality assessment of the total RNA using a NanoDrop and gel electrophoresis indicated that the RNA was of high quality, with a RNA integrity number (RIN) greater than 7. RNA was stored at - 80 degrees Celsius for several days before being shipped for sequencing.

One male and one female 75 bp paired-end sequencing library was prepared using the TruSeq Stranded mRNA Library prep kit and sequenced on an Illumina NextSeq500 at the Ramaciotti Centre for Genomics, University of New South Wales, Australia. In total 79 M and 88 M reads were produced for males and females respectively. Quality assessment of the RNA-seq data using FastQC (Andrews 2010) indicated that the reads were of a high quality and therefore no trimming of reads was performed. The transcriptome was *de novo* assembled for each sex separately using Trinity version 2.1.1 (Grabherr et al. 2011) where all reads were used and the --jaccard_clip option was enabled to minimize gene fusion events caused by UTR overlap in high gene density regions.

### Annotation

Maker version 2.31.8 (Campbell et al. 2014; Holt and Yandell 2011) was used to annotate the PacBio genome via incorporation of *in silico* gene models detected by Augustus (Stanke and Morgenstern 2005) and/or SNAP (Johnson et al. 2008), the *de novo D. serrata* male and female transcriptomes, and protein sequences from 12 *Drosophila* species genomes *(D. ananassae* r1.04, *D. erecta* r1.04, *D. grimshawi* r1.3, *D. melanogaster* r6.07, *D. mojavensis* r1.04, *D. persmillis* r1.3, *D. pseudoobscura pseudoobscura* r3.03, *D. sechellia* 1.3, *D. simulans,* r2.01, *D. virilis* r1.03, *D. willistoni* r1.04, and *D. yakuba* r1.04) obtained from FlyBase (McQuilton et al. 2012; Attrill et al. 2016). Repeat masking was performed based on *D. melanogaster* training (Smit et al. 1996). Maker was run with default settings apart from allowing Maker to take extra steps to identify alternate splice variants and correct for erroneous gene fusion events.

#### Data Availability Statement

All sequence data including PacBio and RNA-seq reads have been submitted to public repositories and are available via the *D. serrata* genome NCBI project accession PRJNA355616. The annotation tracks will be made available in gff formats from www.chenowethlab.org/resources/serratagenome/uponpublication). We also supply a list of *D. melanogaster* orthologs in supplementary file S1.

## RESULTS & DISCUSSION

To assemble a draft *D. serrata* genome we sequenced DNA from a pool of adult males and females that originated from a single inbred line to a coverage of approximately 65x using Pacific Bioscience (PacBio) long-read, single-molecule realtime (SMRT) sequencing technology. We produced 136,119 filtered subreads with a mean read length of 8,940 bp and a read N50 of 12,200 bp that spanned greater than ~13 Gbp (Figure S1). The PacBio reads were assembled using the MHAP algorithm within the Cetera Assembler (Miller et al. 2008; Berlin et al. 2015) after selfcorrection using PBDagCon (Chin et al. 2013). The final genome was polished with a single iteration of Quiver (Chin et al. 2013) and consisted of 1,360 contigs containing more than 198 Mbp with a GC content of 39.13% (Table 1). The longest contig was ~7.3 Mbp and the N50 of all contigs was ~0.95 Mbp. Flow cytometry studies suggest that species of the *montium* subgroup commonly have genome lengths over 200 Mbp (Gregory and Johnston 2008) with the estimate for the female *D. serrata* genome being approximately 215 Mbp (0.22 pg). This estimate is in broad agreement with our assembly length of 198 Mbp for the female genome.

**Table 1:** *D. serrata* genome assembly statistics. Contig length percentages refer to percent total length in each size bin.

| Description |  |
| --- | --- |
| Number of contigs | 1,360 |
| Genome size (bp) | 198,298,763 |
| Longest contig (bp) | 7,300,740 |
| < 1 Kbp | 0.0% |
| 1-10 Kbp | 3.3% |
| 10-100 Kbp | 78.8% |
| 100-1000 Kbp | 15.3% |
| > 1 Mbp | 2.6% |
| N50 (bp) | 942,627 |
| GC content | 39.13% |

### Completeness

Genome completeness was assessed using BUSCO gene set analysis version 2.0 which includes a set of 2799 genes specific to Diptera (Simao et al. 2015). The *D. serrata* assembly contained 96.2% of the BUSCO genes with 94.1% being complete single-copy (defined as complete when the gene’s length is within two standard deviations of the BUSCO group’s mean length) and 2.5% detected as fragmented. Only 1.3% of the BUSCO genes were not found in the assembly (Table 2). In comparison, our analysis of the *D. melanogaster* genome (version r6.05) found it to contain 98.7% complete BUSCO genes. As a further point of comparison we computed BUSCO metrics for a recent PacBio-only assembly of the *D. melanogaster* ISO1 strain genome using all 790 contigs rather than the 132 that were constructed from > 50 reads only (http://www.cbcb.umd.edu/software/PBcR/MHAP/Jquiveredfullassembly!). We also analysed the only other member of the montium subgroup with a publically available genome assembly, *D. kikkawai,* (https://www.hgsc.bcm.edu/arthropods/drosophila-modencode-proiect; NCBI PRJNA62319). Although these assemblies also tended to contain marginally lower numbers of missing BUSCOs, metrics were generally very similar (Table 2) indicating high level of completeness for the *D. serrata* assembly.

**Table 2:** BUSCO gene content assessment for *D. serrata* and two different *D. melanogaster* assemblies, version r6.05 from www.flybase.org. and the full ISO 1 pacbio assembly of Berlin et al. (2015) consisting of 790 contigs. also constructed with the PBcR pipeline. A total of 2799 BUSCOs were searched that form a set of 570 highly conserved Dipteran genes.

|  | *D. serrata* | *D. kikkawai* | *D. melanogaster* |  |
| --- | --- | --- | --- | --- |
| Category |  |  | r6.05 | PacBio |
| Complete BUSCOs: |  |  |  |  |
| Single-copy (%) | 94.1 | 97.1 | 98.2 | 97.7 |
| Duplicated (%) | 2.1 | 1.0 | 0.5 | 0.6 |
| Fragmented BUSCOs (%) | 2.5 | 1.2 | 0.8 | 0.8 |
| Missing BUSCOs (%) | 1.3 | 0.8 | 0.5 | 0.9 |

### Fragmentation and Mis-assemblies

Although our assembly N50 was at the upper end of what might be expected for a short read assembly, it is much lower than a recent PacBio only assembly of the *D. melanogaster* genome (Berlin et al. 2015). There are several reasons why this might be the case. First, we report metrics on all contigs in the assembly rather than excluding those that incorporated fewer than 50 reads as was the case for the *D. melanogaster* assembly (Berlin et al. 2015). Excluding such contigs resulted in an assembly of only 273 contigs with a total genome length of 175 Mbp (vs. 198Mbp) and an N50 of 1.4 Mbp. In this reduced assembly, half of the genome was represented in only 25 contigs, which is closer to the performance seen for *D. melanogaster*. While contigs with less than 50 read support were generally short (median: 23.5 kbp, range: 6.3-110 kbp) and may be excluded in some cases on the basis of quality, when we examined the *D. serrata* annotation data we saw that many of these contigs contained predicted genes that had RNA-seq support, including 14 complete singlecopy BUSCOs. We have therefore retained all contigs in our assembly.

Second, although our N50 filtered subread length of 12,200 kbp is on par with the *D. melanogaster* P5C3 filtered subread lengths (12.2-14.2 kbp) (Kim et al. 2014), we had approximately half the coverage of the *D. melanogaster* assembly (65x vs 130x), which may have reduced our ability to span repetitive regions of the *D. serrata* genome. To examine this further, we reran the PBcR pipeline with *D. melanogaster* data from (Kim et al. 2014) but downsampled it to 65x. We did not see genome contiguity drop to the levels seen for *D. serrata* (data not shown) and note that similar findings were observed by Chakraborty et al. (2016; Figure 5). It therefore seems likely that the *D. serrata* genome, which is longer than that of *D. melanogaster,* may contain longer repetitive regions. Therefore, adequate repeat-spanning coverage would presumably require additional very long reads to achieve the same assembly contiguity seen for *D. melanogaster*. A third factor possibly contributing to a higher degree of fragmentation in our assembly is residual heterozygosity which may have been higher in our *D. serrata* line, which had at least 11 generations of inbreeding, than the ISO1 *D. melanogaster* line.

Because physical maps indicate very strong chromosome arm-level conservation of gene content between *D. serrata* and *D. melanogaster* (Stocker et al. 2012), we examined possible mis-assemblies between chromosomal arms by aligning the six largest contigs (total length ~ 37 Mbp) to the *D. melanogaster* genome using MUMmer (Kurtz et al. 2004). If there were no chromosome arm misplacements, then it was expected that each contig would align to a single *D. melanogaster* chromosome arm, albeit fragmented due to changes in gene order. This was largely the case (Figure 1A), where each contig aligned to a single *D. melanogaster* chromosome arm but with minor sections of alignment to other chromosome arms towards the contig edges where repetitive elements were more likely to be found. The one major exception to this general pattern of conservation was the longest contig in the assembly, contig 3208, that aligned mainly to *D. melanogaster* 3R but contained an approximately 600 kbp segment that aligned to *D. melanogaster* 3L. To test whether this was likely to be a mis-assembly, we searched the contig for previously published SNP markers that have been placed on the *D. serrata* linkage map. The marker m25 (Stocker et al 2012), which maps to 3L, was located in the suspected mis-assembled region (contig 3208, position 3,537,591) indicating that a mis-assembly rather than a genomic translocation rearrangement between 3R and 3L was most likely. The conservation of chromosome arm-level gene content was a common feature of the remaining contigs as well. For instance, while only 354 contigs contained significant tBLASTx hits to at least one *D. melanogaster* gene (genome version 6.05), these contigs spanned 167 Mbp, and the vast majority had greater than 95% tBLASTx hits to a single *D. melanogaster* chromosomal arm (mean = 96.35%, median = 100%) (Figure 2).

**Figure 1.**
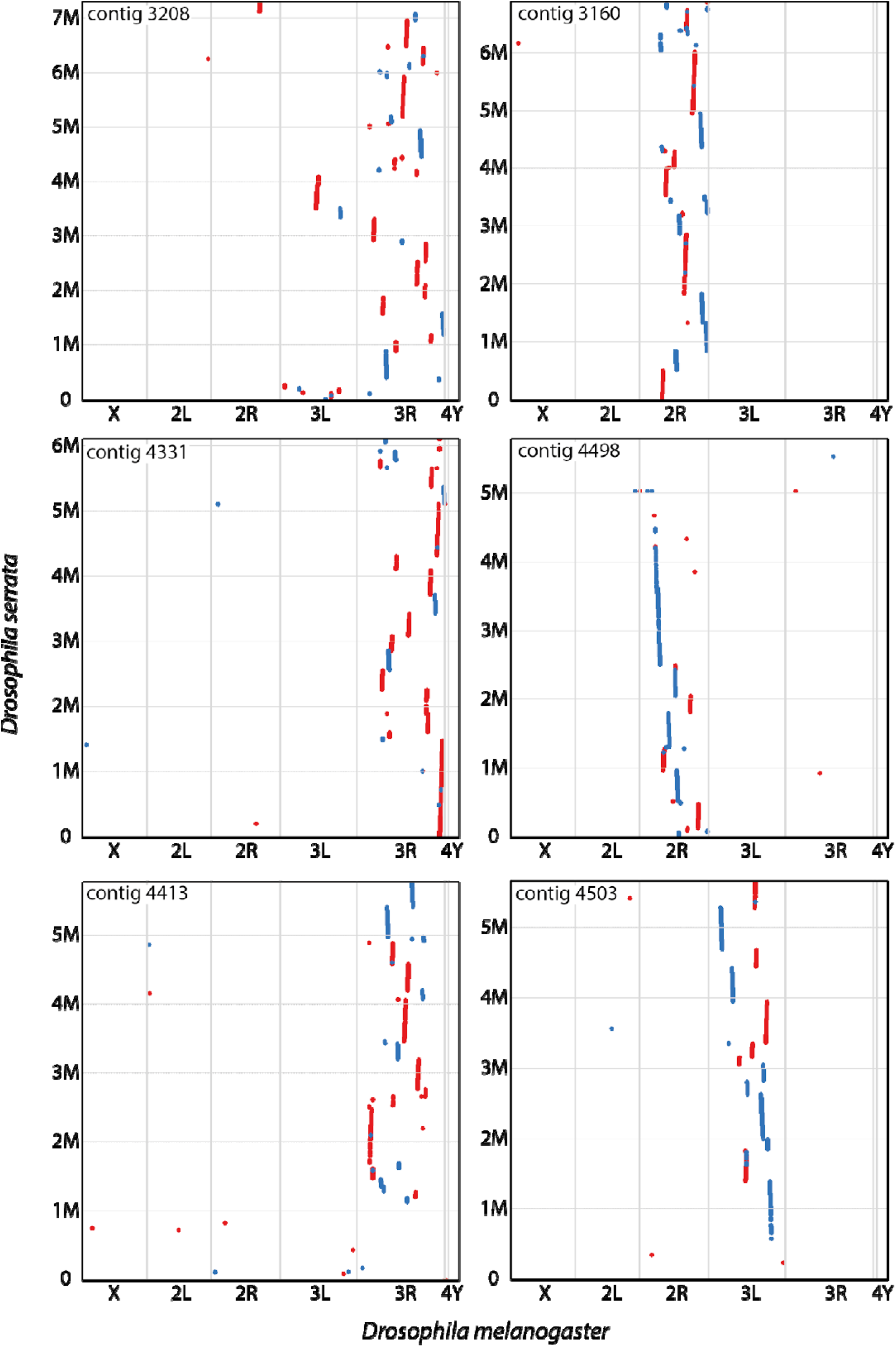
Alignment of the six longest contigs from the *D.serrata* assembly, to the *D. melanogaster* genome version 6.05. Red dots indicate a MUMmer hit that aligns to the *D. melanogaster* genome in the forward orientation; blue dots indicate a MUMmer hit that aligns to the *D. melanogaster* genome in the reverse orientation.

**Figure 2.**
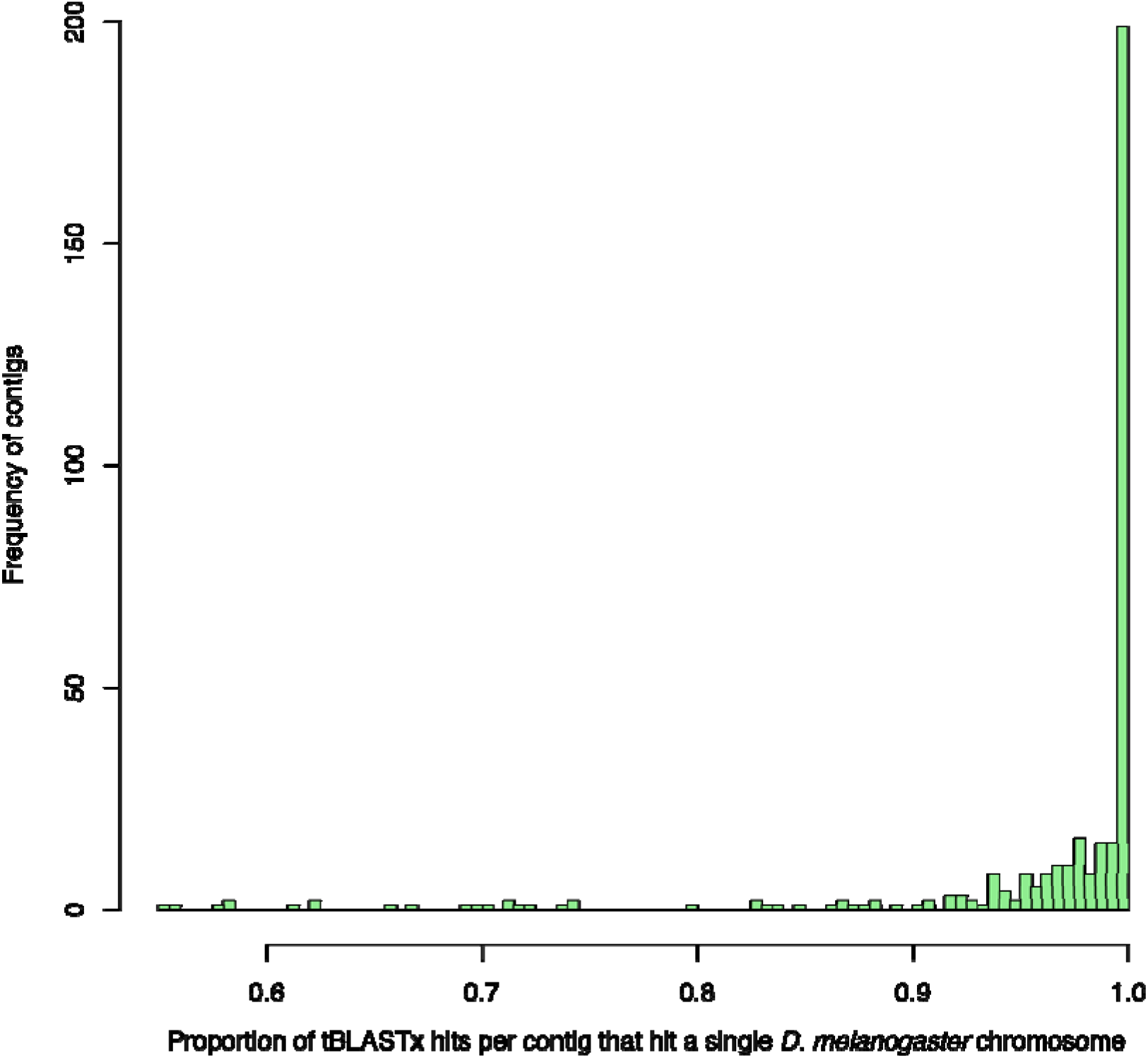
Comparison of *D. serrata* gene locations relative to *D. melanogaster.* On average > 95% of tBLASTx hits to *D. melanogaster* genes (version 6.05) in each contig map to a single *D. melanogaster* chromosome arm.

### Annotation

To facilitate annotation of the *D. serrata* genome we sequenced mRNA from male and female adult flies. The *in silico* gene predictors SNAP (Johnson et al. 2008) and Augustus (Stanke and Morgenstern 2005), found 22,718 and 15,984 genes respectively. Of these *in silico* predicted genes, a total of 14,271 protein coding genes were sufficiently supported by RNA-seq and/or protein sequence data to be annotated by Maker2 (16,202 transcripts) (Holt and Yandell 2011). While the number of genes we annotated in *D. serrata* is similar to the 13,929 protein coding genes that have currently been annotated in *D. melanogaster* (genome version 6.05), we annotated far fewer total transcripts (31,482 identified in *D. melanogaster)* (Attrill et al. 2016), this is likely due to the larger number of tissue types and life stages for which *D. melanogaster* gene expression has been characterized with RNA-seq. Maker scores annotations using the annotation edit distance (AED), a zero-to-one score where a value of zero indicates that the *in silico* annotation and the empirical evidence are in perfect agreement and a value of one indicates that the *in silico* annotation has no support from empirical data (Eilbeck et al. 2009). The AED for the *D. serrata* genome had a mean score of 0.18 and median of 0.13 suggesting that most annotations were of high quality with strong empirical support. Considering that in *Drosophila* appreciable numbers of genes peak in expression during early life stages such as embryogenesis (Arbeitman et al. 2002), our use of adult fly RNA-seq data may mean that some such genes are yet to be annotated. Furthermore, as we used mRNA-seq we have not yet annotated non-coding genes of which there are 3,503 in the *D. melanogaster* genome (Attrill et al. 2016). Future RNA-seq datasets will be used to update the existing gene models.

We observed differences in gene, exon and intron lengths between *D. serrata* and *D. melanogaster*. In *D. serrata* there were on average 3.9 exons per protein coding gene and the gene, exon, and intron lengths were 4,655 bp, 451 bp, and 699 bp respectively. Apart from average exon number which does not differ between the two species, these values are lower than those for *D. melanogaster* protein coding genes (genome version 6.05): where the mean gene, exon, and intron lengths are 6,962 bp, 539 bp, and 1,704 bp respectively (Attrill et al. 2016). The lower average intron length observed in *D. serrata* may be a consequence of annotating far fewer alternate splice variants. In total, coding sequence comprised 33.6% of the genome when including introns and 15.4% of the genome when considering only exons. Lower percentage intron content has been associated with overall longer genomes in the Drosophilidae (Gregory and Johnston 2008) which is consistent with our observations here.

Many of the annotated genes in *D. serrata* were found to be putative orthologs of *D. melanogaster genes* (Supp data file S1). In total 13,141 (92%) were found to be orthologs via best reciprocal BLAST (Huynen and Bork 1998; Moreno-Hagelsieb and Latimer 2008; Tatusov et al. 1997) using tBLASTx with default settings (Camacho et al. 2009) and version 6.05 of the *D. melanogaster* genome (Drosophila 12 Genomes et al. 2007; McQuilton et al. 2012). The median e-value was zero whereas the mean when comparing *D. serrata* genes to *D. melanogaster* was 2.37e^-04^ and when comparing *D. melanogaster* genes to *D. serrata* was 1.55e^-05^, the correlation between e-values for the reciprocal BLAST was 0.88.

### Conclusion

We have assembled a draft genome for a species with no existing genome using only 3GS data. Our study indicates the feasibility of long read-only genome assembly for non-model species with modest sized genomes when using an inbred line. While either greater 3GS coverage or a hybrid merged assembly (Chakraborty et al. 2016) may be required to provide greater genome contiguity, it is clear the genome has a high degree of completeness in terms of gene content and that mis-assemblies at chromosome arm level are rare. The genome and its initial annotation provide a useful resource of future population genomic and trait mapping studies in this species.

## ACKNOWLEDGEMENTS

We thank S. Koren for advice regarding the PBcR pipeline. Funding for this research was provided by The University of Queensland.

---

**Supplementary figure 1.**
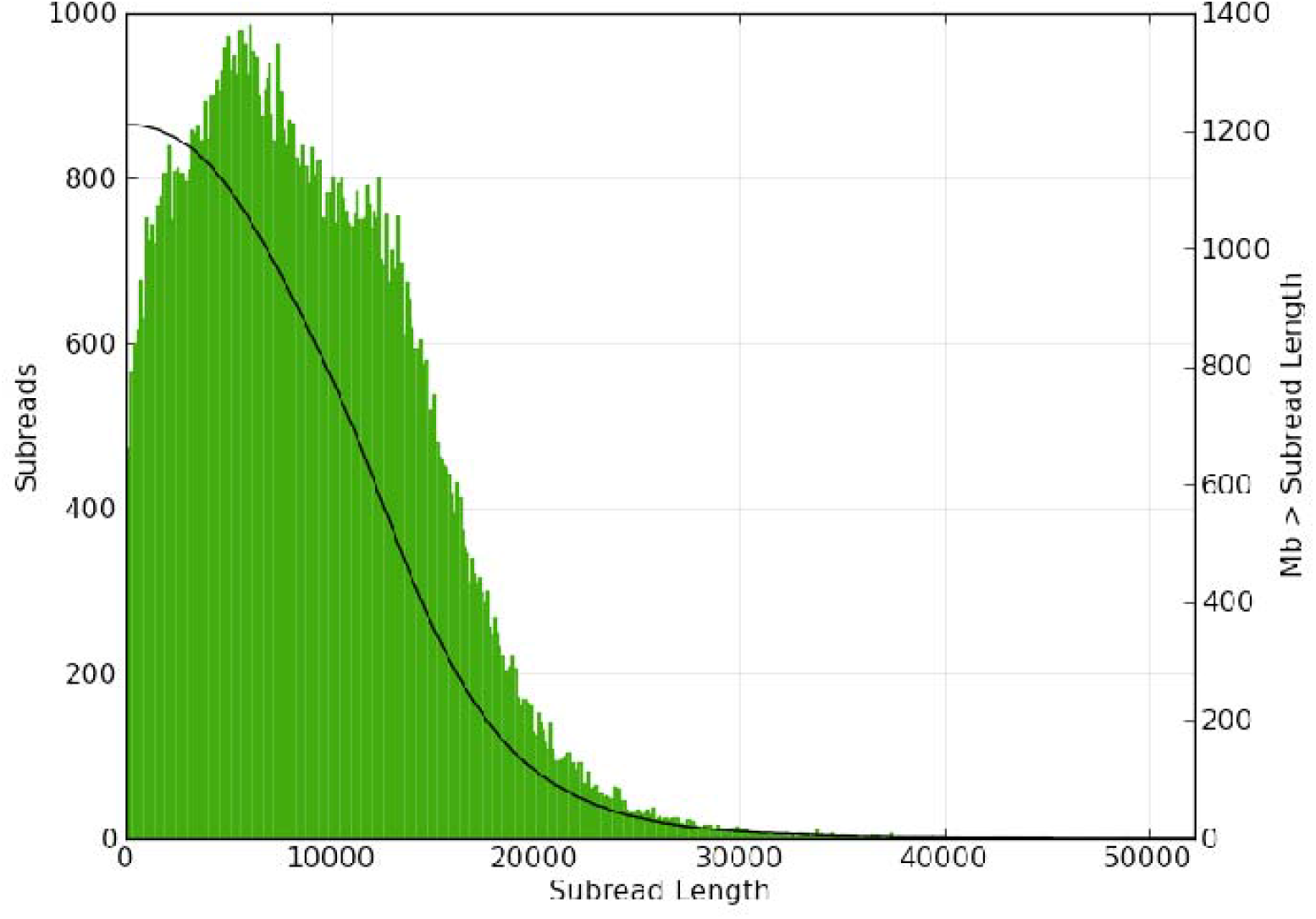
Distribution of filtered subread lengths from 11 SMRT cells 535 on the RS II with P6C4 chemistry.

Supplementary Table 1. List of *D. serrata* to *D. melanogaster* orthologs.

